# Genetic signature of prostate cancer resistant to optimized hK2 targeted alpha-particle therapy

**DOI:** 10.1101/754036

**Authors:** Mesude Bicak, Katharina Lückerath, Teja Kalidindi, Sven-Erik Strand, Michael Morris, Caius Radu, Robert Damoiseaux, Norbert Peekhaus, Austin Ho, Darren Veach, Ann-Christin Malmborg Hager, Steven M Larson, Hans Lilja, Michael R McDevitt, Robert J. Klein, David Ulmert

## Abstract

Hu11B6 is a monoclonal antibody that internalizes in cells expressing androgen receptor (AR)-regulated prostate specific enzyme human kallikrein 2 (hK2; *KLK2*). In multiple rodent models, Actinium-225 labeled hu11B6-IgG_1_ ([^225^Ac]hu11B6-IgG_1_) has shown promising treatment efficacy. In the current study we investigated options to enhance and optimize [^225^Ac]hu11B6 treatment. Firstly, we evaluated the possibility of exploiting IgG_3_, the immunoglobulin G (IgG) subclass with superior activation of complement and ability to mediate FC-gamma-receptor binding, for immunotherapeutically enhanced hK2 targeted alpha-radioimmunotherapy. Secondly, we compared the therapeutic efficacy of a single high activity vs. fractionated activity. Finally, we used RNA sequencing to analyze the genomic signatures of prostate cancer that progressed after targeted alpha therapy. [^225^Ac]hu11B6-IgG_3_ was a functionally enhanced alternative to [^225^Ac]hu11B6-IgG_1_ but offered no improvement of therapeutic efficacy. Progression free survival was slightly increased with a single high activity compared to fractionated activity. Tumor free animals succumbing after treatment revealed no evidence of treatment associated toxicity. In addition to upregulation of canonical aggressive prostate cancer genes, such as *MMP7*, *ETV1*, *NTS* and *SCHLAP1*, we also noted a significant decrease in both *KLK3* (PSA) and *FOLH1* (PSMA) but not in *AR* and *KLK2*, demonstrating efficacy of sequential [^225^Ac]hu11B6 in a mouse model.

## Introduction

Actinium-225 labeled antibodies and small molecules targeting prostate specific antigen (PSMA) are currently being evaluated in clinical trials, and reports have already shown promising therapeutic results in prostate cancer (PCa) patients resistant to androgen receptor (AR)-inhibiting drugs ^1–3^. Unfortunately, toxicities generated by off-target accumulation in radio-sensitive tissues, such as salivary glands and kidneys, is of growing concern ^4,5^. In addition, PSMA expression decreases with increasing AR activity and has been shown to be unevenly expressed in metastases ^6–9^. We have developed an antibody-based platform that targets human kallikrein 2 (hK2), an AR regulated prostate enzyme, specifically expressed at abundant levels in prostate derived tissues. In enzalutamide resistant disease, in which the glucocorticoid receptor (GR) drives the AR-pathway, PSMA expression is lost while hK2 is still highly expressed ^10^. Our group has previously reported on the targeting specificity, efficacy and biological response of radiolabeled hu11B6 as an immunotheranostic vehicle. Preclinical evaluations have been carried out in multiple immunodeficient and immunocompetent, rodent disease models as well as in non-human primates ^3,11–14^. In addition, studies of the uptake mechanism have revealed that hu11B6 is internalized by hK2 expressing cells via a mechanism driven by the neonatal Fc receptor (FcRn); while uncomplexed hu11B6 is released from the cell by recycling endosomes after binding to FcRn, the hu11B6-hK2 complex is routed to lysosomes for processing ^12^. The proximity of the internalized radionuclide-labeled hu11B6 to the nucleus results in efficient cell kill, which is further accelerated by inherent AR and hK2 upregulation related to DNA-repair ^3^.

In the current report we investigated alternative strategies to further improve the therapeutic efficacy of hu11B6. In the field of nuclear medicine, the fragment crystallizable (Fc) is commonly ablated to increase blood clearance. However, these modifications also obliterate possibilities to exploit radiolabeled monoclonal antibodies (mAbs) to orchestrate Fc-receptor activated cytotoxic effects. IgG_3_ activates complement and Fc-gamma-receptor (FcγR)-mediated functions more efficiently than other subclasses. We therefore investigated if changing the subclass of hu11B6 from IgG_1_ to IgG_3_ could improve therapeutic efficacy. However, wild-type IgG_3_ has low FcRn binding capacity due to an arginine at amino acid (aa) position 435 (R435), abrogating cellular internalization ^15^. To overcome this, we constructed a H435 (i.e., histidine at aa 435) containing hu11B6 IgG_3_ allotype, generating hu11B6-IgG_3_^R435H^. As another optimization step, we evaluated therapeutic efficacy of a single high activity vs. two lower activity fractions. Lastly, we examined the genetic profile of PCa tumors resistant to [^225^Ac]hu11B6-IgG_1_ therapy. In summary, while IgG_3_ has shown increased immunotherapeutic efficacy compared to IgG_1_ 16-18, no advantage was shown when applied as a radiolabeled compound targeting hK2. Although susceptible to retreatment, tumors relapsing after [^225^Ac]hu11B6-IgG_1_ showed upregulation of several oncogenes related to aggressive disease.

## Results

### Radiochemistry

The radiochemical yield for the [^225^Ac]DOTA-hu11B6 product using a one-step labeling method was 60.2% ± 29.3% (n=5). The product was 98.5% ± 2.0% (n=5) radiochemically pure and the specific activity was 0.068Ci/g±0.032Ci/g (2.516 GBq/g ± 1.184 GBq/g; n=5). The radiochemical yields for the four hu11B6 combinations prepared using a two-step radiolabeling method were as follows: [^225^Ac]hu11B6-IgG_1_ (native IgG_1_), 3.7% ± 2.1% (n=13); [^225^Ac]hu11B6-IgG_1_H435A (H435A mutant IgG_1_), 3.1% ± 1.0% (n=3); [^225^Ac]hu11B6-IgG_3_ (wild-type IgG_3_), 11.0% (n=1); and [^225^Ac]hu11B6-IgG_3_^R435H^ (R435H mutant IgG), 9.1% (n=1). The radiochemical purity was as follows: native IgG_1_, 99.3% ± 0.51%; H435A mutant IgG_1_, 99.7% ± 0.21%; wild-type IgG_3_, 99.8%; and R435H mutant IgG_3_, 99.8%. The specific activity was as follows: native IgG_1_, 0.079 Ci/g ± 0.055 Ci/g (2.923 GBq/g ± 2.035 GBq/g); H435A mutant IgG_1_, 0.059 Ci/g ± 0.018 Ci/g (2.183 GBq/g ± 0.666 GBq/g); wild-type IgG_3_, 0.068 Ci/g (2.516 GBq/g); and R435H mutant IgG_3_, 0.066 Ci/g (2.442 GBq/g). Comparable radiochemical purity and specific activities were achieved in either process. The KD of [^225^Ac]DOTA-hu11B6, DOTA-hu11B6, and hu11B6 for recombinant hK-2 was 16.7, 16.3, and 15.0 nM, respectively and indicated that the DOTA conjugation and subsequent ^225^Ac-radiolabeling did not affect affinity of the antibody for target analyte.

### The impact of IgG subclass on [^225^Ac]hu11B6 pharmacokinetics

Compared to wildtype [^225^Ac]hu11B6-IgG_3_, significantly (p=0.0139 for all) higher target tissue uptakes were noticed in the GEMM model for FcRn optimized [^225^Ac]hu11B6-IgG_3_^R435H^. In the ventral-, anterior-, and dorsal/lateral prostate lobes of Hi-*Myc* x pb_*KLK2* mice, percent injected activity per gram (%IA/g; ±SD) of [^225^Ac]hu11B6-IgG_3_^R435H^ were 7.75±1.99, 3.60±1.14 and 7.08±3.65, while %IA/g (±SD) [^225^Ac]hu11B6-IgG_3_ were 2.48±0.50, 1.03±0.17 and 0.80±0.26, respectively, at 10d post-injection (Figure 1a). LNCaP-AR tumor models led to similar findings; [^225^Ac]hu11B6-IgG_3_R435H showed significantly (p=0.1653 at 4h, p=0.0003 at 72h, p=0.0132 at 240h p.i.) higher tumor uptake compared to [^225^Ac]hu11B6-IgG_3_, 8.45±4.28 and 0.86±0.79 %IA/g (±SD), respectively, at 10d post-injection (Figure 1b). As expected, enhancement of FcRn binding resulted in a strong trend towards longer blood circulation time of [^225^Ac]hu11B6-IgG_3_^R435H^ (vs. [^225^Ac]hu11B6-IgG_3_) in both genetically modified mouse models (GEMM; p=0.0736 at 240h post injection [p.i.]) and tumor xenograft models (p=0.0013 at 4h, p=0.0552 at 72h, p>0.9999 at 240h p.i.). Compared to the IgG_1_ variant, [^225^Ac]hu11B6-IgG_3_^R435H^ showed significantly lower target tissue accumulation (LNCaP-AR, p=0.0098 and GEMM, p=0.0299 for IgG3^R435H^ at 240h vs. IgG_1_ at 400h p.i.) and shorter retention time in blood (LNCaP-AR, p=0.0056 and GEMM, p=0.4238 for IgG3^R435H^ at 240h vs. IgG_1_ at 400h p.i.) at the final distribution timepoint, while significantly higher (p≤0.0174 for all in both models) accumulation of [^225^Ac]hu11B6-IgG_3_^R435H^ was noted in immunoglobulin degrading organs (i.e., liver and spleen).

**Figure 1.**
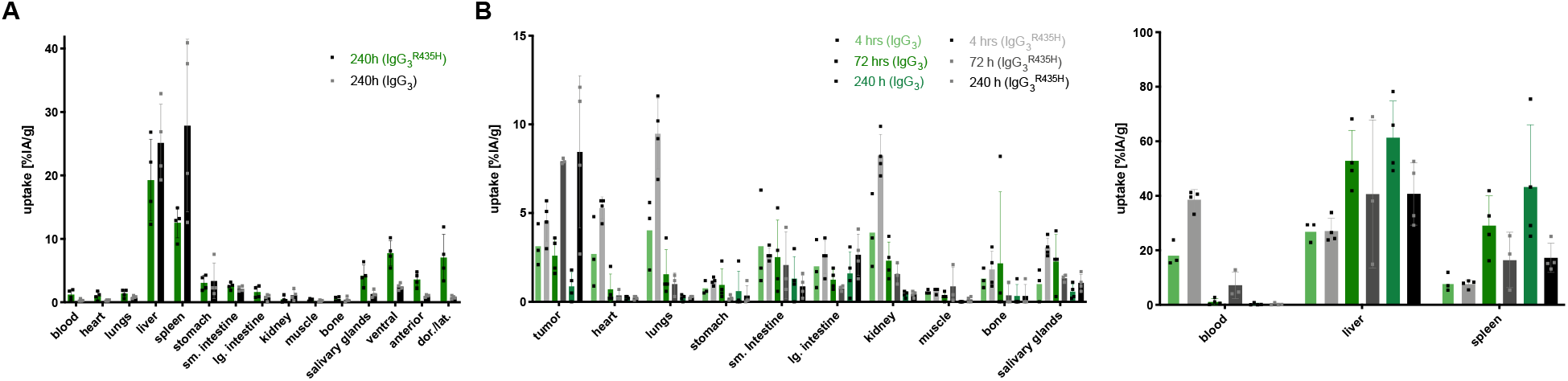
Favorable biodistribution of FcRn optimized [^225^Ac]hu11B6-IgG_3_^R435H^. Mice received 300 nCi (11 kBq) [^225^Ac]hu11B6-IgG_3_ or [^225^Ac]hu11B6-IgG_3_^R435H^. Organs were collected at indicated time points (h) and specific uptake (%IA/g) was determined with a gamma-counter. **(A)** Hi-Myc_*KLK2* GEMM. **(B)** LNCaP-AR xenografts. For better visualization, biodistribution data are shown in two separate graphs with different y-axis scales. Columns and bars represent mean±SD (n=5 mice/group). Individual values are shown as squares. [^225^Ac]hu11B6-IgG_3_ treatment: light green - 4 h; middle green - 48 h; dark green - 240 h. [^225^Ac]hu11B6-IgG_3_^R435H^: light grey - 4 h; middle grey - 48 h; dark grey - 240 h. d*or./lat. - dorsal and lateral prostate; lge. int. - large intestine; sm. intestine - small intestine*

### The impact of IgG subclass on [^225^Ac]hu11B6 alpha-radioimmunotherapy (RIT) in LNCaP-AR s.c. xenografts

Mice with LNCaP-AR s.c. xenografts were randomized to receive (A) [^225^Ac]hu11B6-IgG_1_ (n=5), (B) [^225^Ac]hu11B6-IgG_1_^H435A^ (IgG_1_ with inhibited FcRn-coupled cellular processes by exchanging histidine 435 to an alanine) (n=5), (C) [^225^Ac]hu11B6-IgG_3_^R435H^ (n=5) or (D) [^225^Ac]hu11B6-IgG_3_ (n=5). [^225^Ac]hu11B6 constructs based on FcRn complexing Ig-subforms resulted in significantly better treatment outcomes (Figure 2, **Supplemental Figure 1**). Kaplan– Meier analysis of the four treatment groups showed median survival of 18.43 (A), 9.14 (B), 11.42 (C) and 7.86 (D) weeks after administration of 300 nCi (11 kBq), respectively (A vs. B, p<0.0001; C vs. D, p=0.0020; A vs. C, p=0.0536) (Figure 2a). Assessment of volumetric effects showed that mice treated with [^225^Ac]hu11B6-IgG_3_^R435H^ had reduced tumor growth kinetics compared to [^225^Ac]hu11B6-IgG_3_ (Figure 2b). However, [^225^Ac]hu11B6-IgG_1_ outperformed [^225^Ac]hu11B6-IgG_3_^R435H^ (Figure 2c); at the conclusion of the study, half of the animals treated with the specific and internalizing [^225^Ac]hu11B6-IgG_1_ drug were alive, while only one animal was alive in the group that received [^225^Ac]hu11B6-IgG_3_^R435H^.

**Figure 2.**
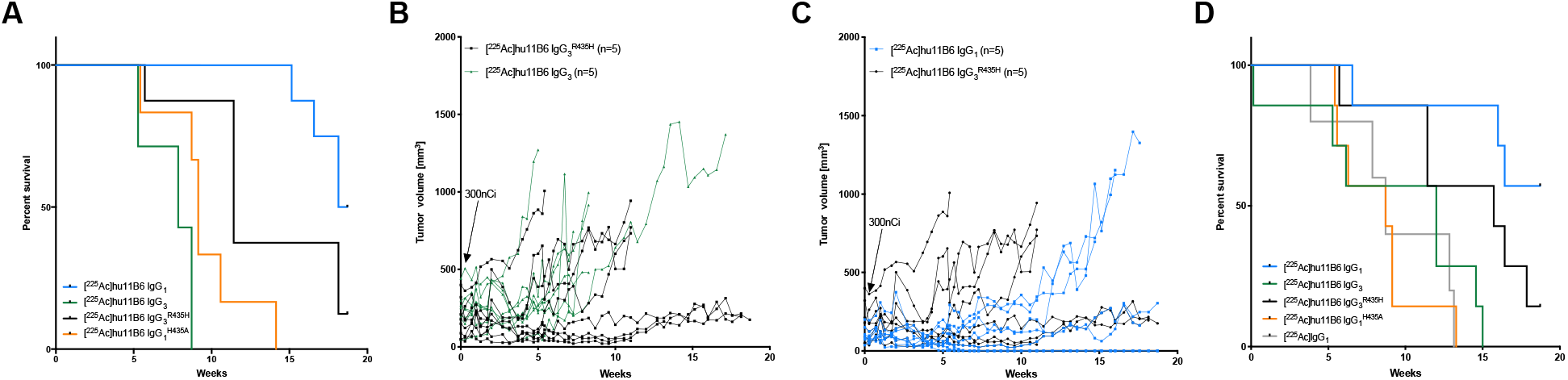
The impact of IgG subclass on [^225^Ac]hu11B6 alpha-radioimmunotherapy. **(A-C)** Mice with LNCaP-AR s.c. xenografts were randomized to receive 300 nCi (11 kBq) [^225^Ac]hu11B6-IgG_1_ (n=5; blue),[^225^Ac]hu11B6-IgG_1_^H435A^ (n=5; orange), [^225^Ac]hu11B6-IgG_3_^R435H^ (n=5; black) or [^225^Ac]hu11B6-IgG_3_ (n=5; green). (**A**) Kaplan-Meier curves for survival. (**B**) Tumor volumes in individual mice following treatment with IgG_3_ or IgG_3_^R435H^ antibodies, respectively. (**C**) Tumor volumes in individual mice following treatment with IgG_1_ or IgG_1_^H435A^ antibodies, respectively. (**D**) Kaplan-Meier curves for survival of Hi-Myc_*KLK2* GEMM following treatment with 300 nCi (11 kBq) [^225^Ac]hu11B6-IgG_1_ (n=7; blue), [^225^Ac]hu11B6-IgG_1_^H435A^ (n=7; orange), [^225^Ac]hu11B6-IgG_3_^R435H^ (n=7; black), [^225^Ac]hu11B6-IgG_3_ (n=7; green) or non-specific [^225^Ac]IgG_1_ (n=5; grey).

### The impact of IgG subclass on [^225^Ac]hu11B6 alpha-RIT in immunocompetent PCa GEMM

40-55 week old hK2-expressing Hi-Myc GEMM were randomized to receive (A) [^225^Ac]hu11B6-IgG_1_ (n=7), (B) [^225^Ac]hu11B6-IgG_1_^H435A^ (n=7), (C) [^225^Ac]hu11B6-IgG_3_^R435H^ (n=7), (D) [^225^Ac]hu11B6-IgG_3_ (n=7) or (E) non-specific [^225^Ac]IgG_1_ (n=5). Kaplan–Meier analysis of these four treatments showed median survival of undefined (A; >50% of the cohort had survived with [^225^Ac]hu11B6-IgG_1_), 8.71 (B), 15.71 (C), 12.0 (D) and 8.72 (E) weeks after administration of 300 nCi (11 kBq), respectively (A vs. B, p=0.0020; C vs. D, p=0.0576; A vs. C, p=0.1197) (Figure 2d). At the end of the study, 4 of 7 (57%) GEMM were alive in group A but only 1 of 7 (14%) in group C. These data indicate that an IgG_1_ based hK2 targeting compound is more effective than IgG_3_R435H when applied for alpha-particle RIT. Therapeutic effect by ineffective FcRn-binding compounds (B, D) were not significantly different from non-specific [^225^Ac]IgG_1_ (B vs E, p=0.9738; D vs E, p=0.5726), showing that efficacy is dependent on a combination of specific targeting and cellular internalization.

### Pharmacokinetics of [^225^Ac]hu11B6-IgG_1_, labeled using a novel 1-step protocol, in s.c. PCa xenografts and immunocompetent PCa GEMM

The one-step labelling protocol based on pre-conjugated hu11B6 was developed to facilitate clinical translation. Biodistribution of [^225^Ac]hu11B6 was studied in LNCaP-AR (*KLK2*+) s.c. xenografts at 48, 120 and 400 h p.i., in PC3 (*KLK2*-) s.c. xenografts at 400 hrs p.i. and in 15-30 weeks old hK2-expressing Hi-Myc GEMM at 400 hrs p.i. As expected, uptake of [^225^Ac]hu11B6 increased over time in LNCaP-AR tumors and uptake at 400 hrs p.i. was significantly (p=0.0020) higher than in PC3 tumors at the same time-point (Figure 3a). In GEMM, the highest specific uptake of [^225^Ac]hu11B6 was found in prostate lobes (Figure 3b). In summary, biodistribution profile of 1-step labeled [^225^Ac]hu11B6 closely resembled our previously published results based on the 2-step labeled [^225^Ac]hu11B6 compound ^3^. Therefore, the one-step labelling protocol was used for all further studies.

**Figure 3.**
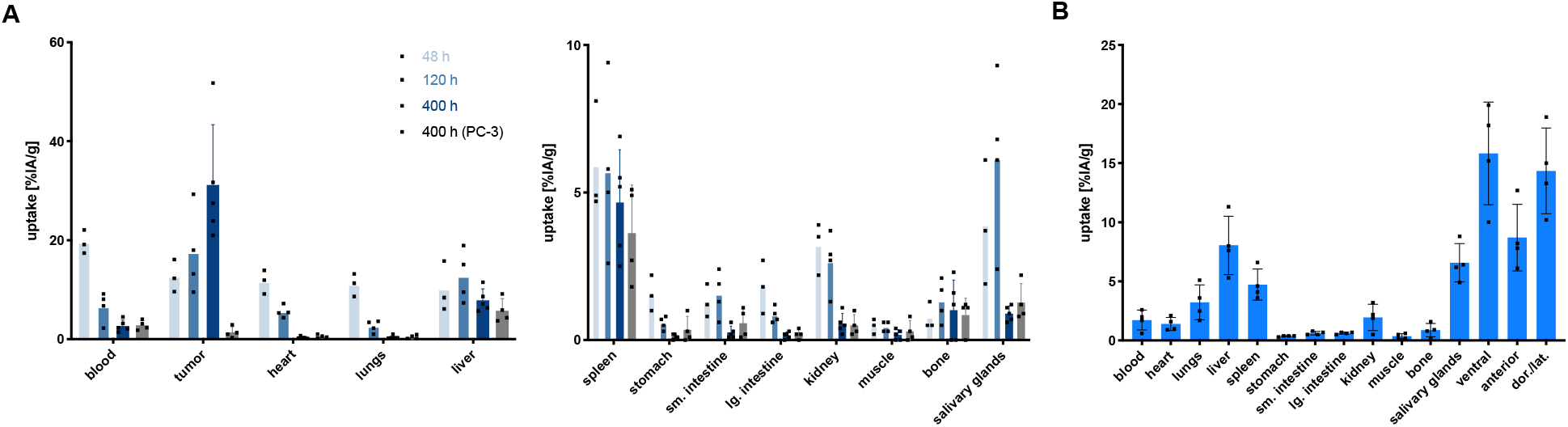
Biodistribution of [^225^Ac]hu11B6-IgG_1_, labeled using a 1-step protocol. Mice received 300 nCi (11 kBq) [^225^Ac]hu11B6-IgG_1_. Organs were collected at indicated time points and specific uptake (%IA/g) was determined in a gamma-counter. **(A)** LNCaP-AR and PC-3 xenografts. LNCaP-AR are shown in blue: light blue - 48 h; middle blue - 120 h; dark blue - 400 h. PC-3 are shown in grey (400 h). For better visualization, biodistribution data are shown in two separate graphs with different y-axis scales. **(B)** Hi-Myc_*KLK2* GEMM, 400 h time point. Columns and bars represent the mean±SD (n=5 mice/group). Individual values are shown as black squares. Time indicates time post radioligand injection. d*or./lat. - dorsal and lateral prostate; lge. int. - large intestine; sm. intestine - small intestine*

### Therapeutic efficacy of fractionated vs. single activity [^225^Ac]hu11B6-IgG_1_ alpha RIT

Bilaterally inoculated LNCaP-AR s.c. xenografts were used in both treatment studies. Therapeutic effects were monitored by tumor size (caliper measurements) and assessment of PSA in blood. In the fractionated treatment experiment, animals (n=10) were randomized to receive 300 nCi (11 kBq) [^225^Ac]hu11B6 (n=5) or no treatment (n=5). Previously treated mice were given a second injected activity (300 nCi; 11 kBq) 20 weeks after the first treatment. Tumors were either fully eradicated or substantially reduced after each treatment round; 5 of 8 (63%) and 2 of 5 (40%) tumors relapsed after the first activity and second activity administration, respectively (Figure 4a; **Supplemental Figure 2a**). In the single high-activity treatment experiment, xenografts (n=10) were randomized to receive a single activity 600 nCi (22 kBq) of [^225^Ac]hu11B6 (n=5) or no treatment (n=5). All tumors were eradicated (tumors not palpable and PSA not detectable); during the study period, only 1 of 6 (17%) tumors relapsed (Figure 4b; **Supplemental Figure 2b**). Mice that had lost weight (≥20% weight loss) at the end of the high-activity study were euthanized and send for autopsy evaluation. These reports concluded age-related macroscopic and histopathologic but no treatment-related pathological changes. Comparing fractionated vs. single activity treatment median survival was not reached (80% of the cohort had survived at the conclusion of the study) following 600 nCi (22 kBq), whereas 2x 300 nCi (2x 11 kBq) resulted in a median survival of 40.3 weeks after the first treatment (p=0.3186 vs. high-activity) (Figure 4c, d).

**Figure 4.**
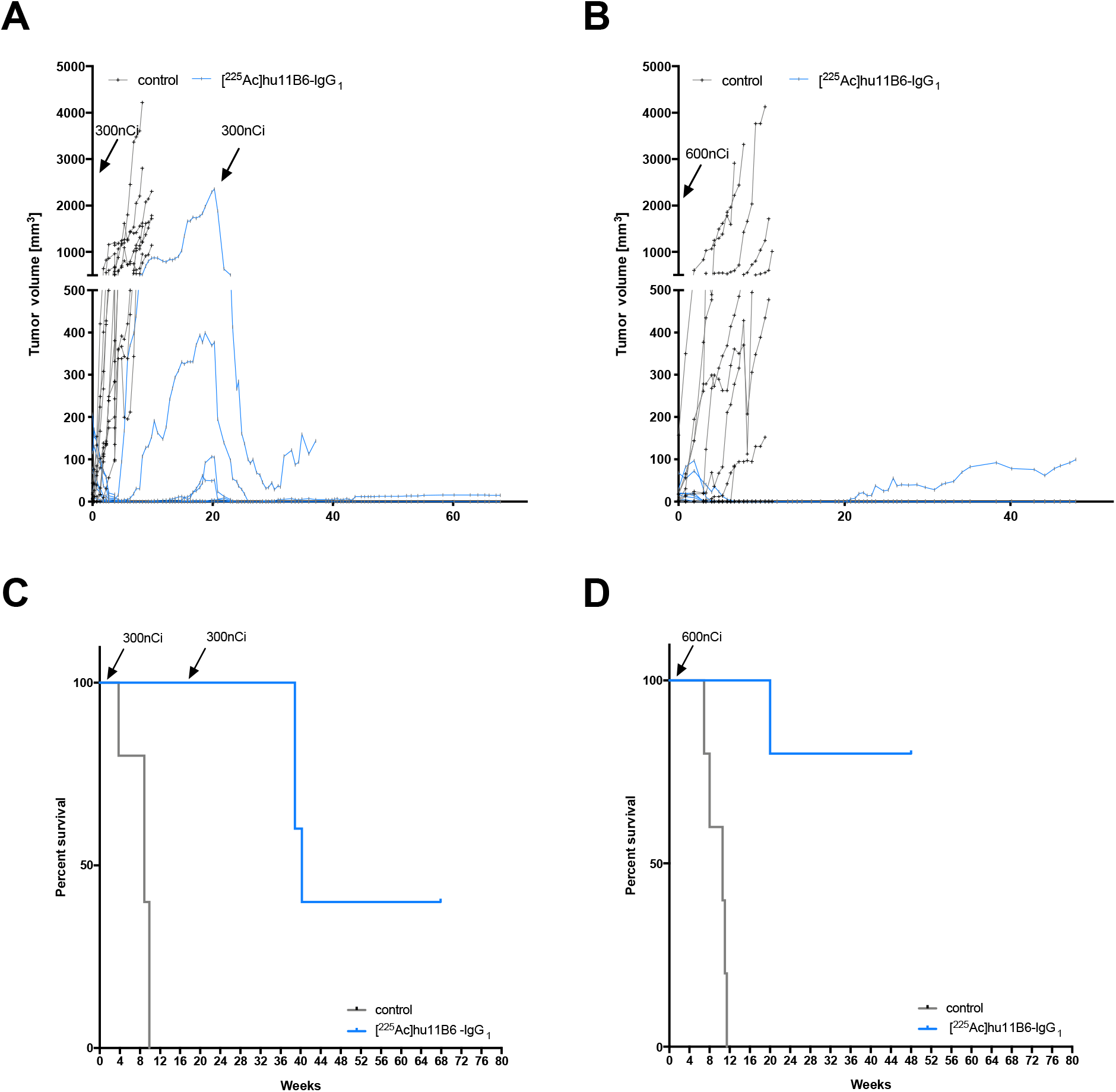
Therapeutic efficacy of fractionated vs. single high-dose [^225^Ac]hu11B6-IgG_1_ alpha-radioimmunotherapy. Mice with LNCaP-AR s.c. xenografts (n=5 mice/group) received 2x 300 nCi (2x 11 kBq; ~4.5 month apart) (**A, C**) or 1x 600 nCi (22 kBq) (**B, D**) [^225^Ac]hu11B6-IgG_1_ (blue; vs. untreated, black). (**A, B**) Tumor volumes of treated and control mice. (**C, D**) Kaplan Meier analysis of survival of treated and control mice.

### Transcriptome Sequencing

The transcriptome of tumors relapsing after 2 cycles [^225^Ac]hu11B6 (vs. untreated control tumors, n=3/group) was profiled. RNA-sequencing identified 58,828 genes of which 1,644 (2.8%; 512 up-, 1,132 down-regulated) were differentially expressed in relapsing tumors. Among the differentially expressed genes (DEG), genes associated with aggressive prostate cancer, such as *MMP7 (matrix metalloprotease 7)*, *ETV1 (ETS translocation variant 1)*, *NTS (neurotensin), TMEFF2* (*transmembrane protein with EGF-like and two follistatin-like domains 2*) and *SCHLAP1 (SWI/SNF complex antagonist associated with prostate cancer 1)*, and androgen repressed genes, such as *PMP22 (peripheral myelin protein 22)*, *CAMK2N1 (calcium/calmodulin dependent protein kinase II inhibitor 1)* and *UGT2B17UDP (glucuronosyltransferase family 2 member B17)*, were significantly up-regulated. In contrast, several AR-regulated genes, e.g. *KLK3*, *FOLH1*, *PMEPA1 (prostate transmembrane protein, androgen induced 1)* and *SPOCK1 (Testican-1)*, were downregulated or remained unchanged (*KLK2, TMPRSS2 [transmembrane serine protease 2], LONRF1 [LON peptidase N-terminal domain and RING finger protein 1], KCNN2 [small conductance calcium-activated potassium channel protein 2]*). Expression of *AR* itself was comparable in treated and untreated tumors (Table 1). Figure 5 gives an overview of all DEGs and Tabels 2 and 3 summarize the top 10 up-regulated and down-regulated genes.

**Table 1.** Differentially regulated genes associated with PCa and AR signaling.

| Gene Name | Fold Change (Log 2) | P-Value | Adjusted P-Value |
| --- | --- | --- | --- |
| **AR** | -0.16 | 0.3125 | 0.89 |
| **canonical aggressive PCa genes** |  |  |  |
| **MMP7** | 7.86 | 0.00E+00 | 0.00E+00 |
| **ETV1** | 10.58 | 0.00E+00 | 0.00E+00 |
| **NTS** | 11.76 | 0.00E+00 | 0.00E+00 |
| **SCHLAP1** | -12.18 | 3.20x10<sup>-150</sup> | 3.39x 10<sup>-147</sup> |
| **Androgen-repressed genes** |  |  |  |
| **CAMK2N1** | -6.24 | 4.17x 10<sup>-150</sup> | 4.22x 10<sup>-147</sup> |
| **UGT2B17** | -3.80 | 4.67x 10<sup>-45</sup> | 5.43x 10<sup>-43</sup> |
| **PMP22** | 1.57 | 1.40x 10<sup>-5</sup> | 8.96x 10<sup>-5</sup> |
| **AR-regulated genes** |  |  |  |
| **TMEFF2** | 7.32 | 1.08x 10<sup>-273</sup> | 4.00x 10<sup>-234</sup> |
| **PMEPA1** | -1.54 | 2.34x 10<sup>-20</sup> | 2.35x 10<sup>-18</sup> |
| **KLK3** | -1.68 | 1.95x 10<sup>-16</sup> | 1.46x 10<sup>-14</sup> |
| **FOLH1** | -0.82 | 4.00x 10<sup>-7</sup> | 9.85x 10<sup>-6</sup> |
| **SPOCK1** | 0.82 | 2.44x 10<sup>-4</sup> | 1.17x 10<sup>-3</sup> |
| **KLK2** | -0.44 | 0.0183 | 0.12 |
| **TMPRSS2** | -0.10 | 0.5442 | 1.00 |
| **LONRF1** | 0.21 | 0.3803 | 0.5191 |
| **KCNN2** | 0.16 | 0.2955 | 0.4319 |

**Table 2.** Top 10 up-regulated genes at relapse after treatment with [^225^Ac]hu11B6-IgG_1_.

| **Gene Name** | **Fold Change (Log 2)** | **P-Value** | **Adjusted P-Value** |
| --- | --- | --- | --- |
| **MMP7** | 7.86 | 0.00E+00 | 0.00E+00 |
| **NCAM2** | 10.54 | 0.00E+00 | 0.00E+00 |
| **ETV1** | 10.58 | 0.00E+00 | 0.00E+00 |
| **NTS** | 11.76 | 0.00E+00 | 0.00E+00 |
| **TMEFF2** | 7.32 | 1.08x 10<sup>-273</sup> | 4.00x 10<sup>-234</sup> |
| **PLA2G2A** | 10.79 | 3.46x 10<sup>-227</sup> | 9.62x 10<sup>-224</sup> |
| **WBP5** | 9.13 | 1.01x 10<sup>-210</sup> | 2.49x 10<sup>-207</sup> |
| **TMEM45B** | 6.39 | 1.78x 10<sup>-208</sup> | 3.96x 10<sup>-205</sup> |
| **CFLAR** | 7.47 | 6.56x 10<sup>-182</sup> | 1.22x 10<sup>-178</sup> |
| **PEG3** | 8.40 | 2.22x 10<sup>-175</sup> | 3.80x 10<sup>-172</sup> |

**Table 3.** Top 10 down-regulated genes at relapse after treatment with [^225^Ac]hu11B6-IgG_1_.

| **Gene Name** | **Fold Change (Log 2)** | **P-Value** | **Adjusted P-Value** |
| --- | --- | --- | --- |
| **LAMC3** | -8.12 | 8.02x 10<sup>-232</sup> | 2.55x 10<sup>-228</sup> |
| **NES** | -6.743 | 1.08x 10<sup>-205</sup> | 2.19x 10<sup>-202</sup> |
| **PASD1** | -11.43 | 4.53x 10<sup>-155</sup> | 5.30x 10<sup>-152</sup> |
| **DENND2D** | -7.54 | 3.34x 10<sup>-153</sup> | 3.71x 10<sup>-150</sup> |
| **NRXN1** | -6.45 | 1.60x 10<sup>-142</sup> | 1.31x 10<sup>-139</sup> |
| **KCNJ3** | -4.73 | 5.93x 10<sup>-136</sup> | 4.39x 10<sup>-133</sup> |
| **PTGFR** | -6.38 | 1.50x 10<sup>-132</sup> | 1.07x 10<sup>-129</sup> |
| **SLC25A43** | -10.94 | 2.87x 10<sup>-122</sup> | 1.72x 10<sup>-119</sup> |
| **FAM213A** | -4.92 | 2.97x 10<sup>-119</sup> | 1.74x 10<sup>-116</sup> |
| **SLC7A5** | -4.24 | 5.65x 10<sup>-114</sup> | 3.14x 10<sup>-111</sup> |

**Figure 5.**
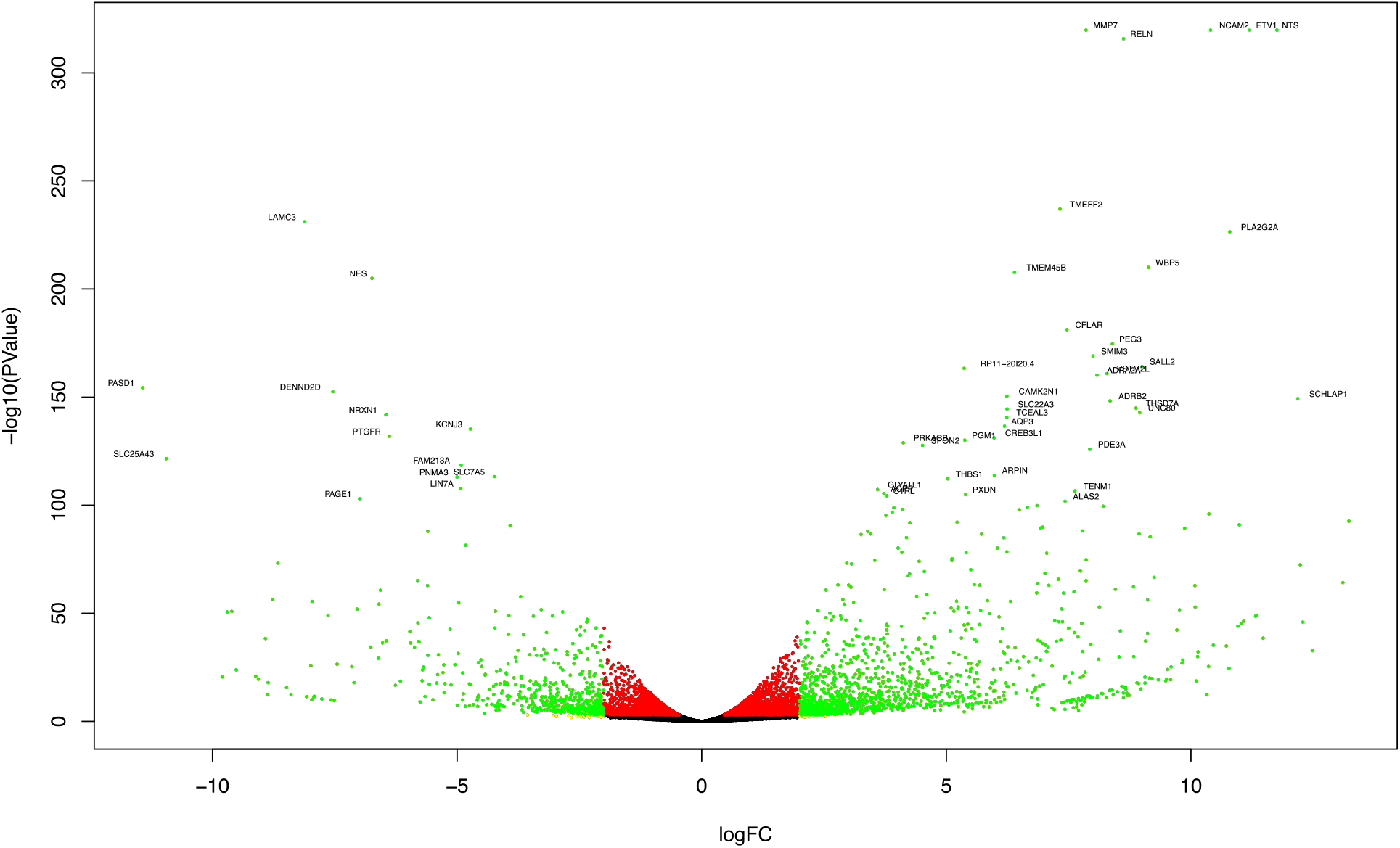
Summary of the RNA-sequencing results by volcano plot representation of differentially expressed genes (DEG). The x-axis shows log2 fold-changes in expression and the y-axis the log10 p-values of a gene being differentially expressed. Top 50 DEGs are labeled. Green dots represent DEGs with p<0.001 and absolute value of log2 fold change >2; yellow dots represent those with only absolute value of log2 fold change >2; red dots represent those with only p<0.001.

## Discussion

Recently approved drugs that target AR signaling such as abiraterone and enzalutamide have rapidly become standard therapies for advanced-stage prostate cancer ^19,20^. Initial responses are usually observed, but in an absolute majority of these patients, reactivation of the AR-pathway inevitably occurs within 6 to 12 months. An alternative treatment option is to target tumors using radiolabeled antibodies or small molecules, a technology that is highly dependent on the specificity of the vehicle and target expression in non-malignant tissues. We have developed an antibody-based platform that targets hK2, an enzyme that is prostate tissue-specific and under direct regulation of AR, the central driver of prostate adenocarcinoma. Our lead antibody, hu11B6, has been thoroughly evaluated for tumor detection and monitoring of disease activity by positron emission tomography and single photon emission computed tomography ^12,14^. We have reported that hu11B6 can be applied for RIT applications ^3,11,13^. In the current report we aimed to evaluate previously untested and innovative methods that could further increase efficacy. First, we evaluated if therapeutic efficacy could be improved by altering the Fc-subclass. Second, we assessed the impact of dosing strategies. Finally, to determine radio-biological mechanisms that could appropriate combination therapies or elucidate potential drug resistance, we genetically profiled tumors that relapsed after prolonged treatment.

Induction of DNA damage, tumor cell death and modulation of tumor microenvironment are the main effects of ionizing radiation to reduce tumor masses, but also to effectuate the immune system ^21^. Therefore, we investigated if the therapeutic efficacy of [^225^Ac]hu11B6 could be improved by exploiting the complement-, and FcγR binding capacities of the mAb. We constructed an Fc-substituted hu11B6; from IgG_1_-Fc to IgG_3_-Fc. Compared to IgG_1_, wild-type IgG_3_ has an arginine instead of a histidine at the aa position 435. Although both residues are positively charged, arginine does not deprotonate at neutral pH, resulting in a less pH dependent binding to FcRn. To rescue FcRn-mediated internalization, we introduced a histidine at the aa position 435 ^22^, generating a hu11B6-IgG_3_-R435H. Indeed, this optimization is critical as almost no effect was noted with [^225^Ac]hu11B6-IgG_3_, while [^225^Ac]hu11B6-IgG_3_-R435H produced a relatively potent therapeutic outcome. Biodistribution in immunodeficient tumor models and immunocompetent GEMM indicated that [^225^Ac]hu11B6-IgG_3_-R435H is less stable than its IgG_1_ counterpart; significantly higher uptake in immunoglobulin-degrading organs were noted. This is probably due to the long hinge region of IgG_3_, which is more prone to proteolytic degradation^23^.

Based on our results, [^225^Ac]hu11B6-IgG_3_-R435H did not show any therapeutic advantage over [^225^Ac]hu11B6-IgG_1_. However, the lack of increased efficacy could be due to critical differences between mouse and human FcγR networks. For example, mouse natural killer (NK) cells express mFcγRIII, while human NK cells express hFcγRIIIA. hFcγRIIIA is a key mediator of antibody therapy in humans, while mFcγRIII only modestly contributes to mAb efficacy ^24,25^. Therefore, experiments based on GEMM expressing human FcγRs would be needed in order to fully elucidate whether RIT efficacy can be enhanced by using Fc-engineered mAbs. Our approach of exploiting Fc-engineered mAbs for RIT is novel. In the field of nuclear medicine, the Fc region has been viewed as redundant and various Fc-edited antibody-like platforms have been invented ^26^. Omitting the Fc-region decreases the half-life of the antibody by escaping FcRn-interaction and permitting renal clearance. However, Fc contains motifs for activation of both effector immune cells and the classical pathway of the complement (C1q). One of the most important breakthroughs in the field of immunology during the last two decades is the discovery and characterization of the Fc-portion interaction with FcγR which orchestrates mAb dependent cellular cytotoxicity through cytotoxic effects via agonistic and antagonistic binding ^27^. The specific FcγR engaged by a given Fc domain is dictated by Fc structure which is determined by the IgG subclass. An example of the multifaceted therapeutic activity are checkpoint-inhibitor mAbs (e.g., anti-CTLA4) that mediate antitumor activity by altering the composition and functional activity of leukocytes within the tumor microenvironment. However, the therapeutic action is also an effect of the capacity of the mAb to activate FcγR expressing macrophages within the tumor microenvironment ^28^. Radiation has been shown to induce immune-modulatory effects. Both preclinical and clinical studies have shown that radiation enhances the activity of immune checkpoint blockers and thereby promotes systemic antitumor responses ^29^. The capacity of radiation therapy to promote immune-modulatory effects is now the subject of intense scientific and clinical investigation. Based on these observations, a radiolabeled full sized mAb might have the unique capacity to target its antigen while inducing immune-modulatory effects and activate Fc receptor mediated cytotoxicity.

Although a single high activity may deliver optimal absorbed dose to kill a larger fraction of tumor cells, fractionated therapy could offer the advantage of lower myelotoxicity and prolonged tumor response. In accordance to previous studies based on [^177^Lu]hu11B6, results showed that the therapeutic effect of [^225^Ac]hu11B6 is absorbed dose dependent; fewer tumors relapsed in mice treated with a higher activity ^13,30^. All of our previous evaluations of [^225^Ac]hu11B6-RIT studies were based on evaluation of a single activity. In the current study, [^225^Ac]hu11B6 treatment was repeated when several mice had relapsed, and relapsed tumors had progressed to a significant volume. Based on tumor and PSA measurements, we did not notice any signs of resistance to treatment when repeated. To retrieve more biologically relevant data, the transcriptome of tumors progressing after [^225^Ac]hu11B6 were evaluated by RNA-sequencing. Although the tumor material post-treatment was limited since only few tumors relapsed and analysis was only based on a single tumor model, the material displays a unique profile. We have previously reported that early effects (400 h after treatment initiation) of [^225^Ac]hu11B6 RIT include DNA-repair associated AR-induction similar to external beam therapy which resulted in increased expression of hK2 and PSA, while PSMA decreased ^3^. In the current study, we expanded these finding to tumors relapsing several months after [^225^Ac]hu11B6 treatment. Compared to treatment naïve tumors, relapsing tumors harbored an altered expression profile of downstream AR-governed biomarkers. Although AR expression remained intact, some of the key AR governed genes (*FOLH1*, *KLK3*, *PMEPA1* and *SPOCK1*) were down-regulated while others (*KLK2* and *TMPRSS2*, *LONRF1, KCNN2*) remained unchanged. Conversely, some of the androgen repressed genes were upregulated (*PMP22*, *CAMK2N1* and *UGT2B17*). Interestingly, these findings bear a resemblance to the genetic changes reported in glucocorticoid receptor driven enzalutamide resistance ^10^. While the underlying radiobiological mechanisms instigating PSMA down-regulation during treatment and in recurrent tumors could be different, these observations could potentially explain the reduced treatment efficacy noted upon repeated PSMA targeted RIT and radioligant therapy ^6,31,32^. Although preclinical models have limitations, the current outcome demonstrates efficacy of serial [^225^Ac]hu11B6 treatment in a mouse model; relapsing tumors in mice express abundant levels of hK2 (*KLK2*) (**Supplemental Figure 2**) and were successfully treated by a second activity. Also, more than a year after treatment no imminent organ toxicities were noted at necropsy.

In addition to deregulated AR regulated genes, further analysis of the transcriptome of relapsing tumors revealed an intriguing profile. Among the most strongly upregulated genes were *ETV1, MMP7, TMEFF2, SCHLAP1 and NTS. ETV1* predisposes prostate cells for cooperation with other oncogenic events such as PTEN loss, leading to more aggressive disease in murine models and human patients ^33^. *ETV1* expression is associated with enrichment of steroid hormone biosynthesis pathway and androgen and estrogen metabolism by binding to *HSD17B7*, a gene shared by steroid biosynthesis and steroid hormone biosynthesis pathways. In accordance with this, we found *HSD17B7* to be significantly up-regulated in [^225^Ac]hu11B6 resistant tumors (p=2.27*10^−7^). *MMP7* is a tumor biomarker associated with increased risk for metastases and poor survival, including prostate cancer ^34,35^. *TMEFF2* has been reported as an AR-regulated protein ^36^ that can function both as an oncogene and as a tumor suppressor by affecting Akt and ERG activation ^37^. While the full length intracellular TMEFF2 acts as a tumor suppressor ^38–40^, the shed ectodomain promotes growth ^39^. Successful immuno-targeting of TMEFF2 expressing PCa tumors has been reported and applied for both positron emission tomography (PET)-imaging and antibody-drug conjugates ^41–43^. Therefore, *in vivo* imaging and co-treatment with TMEFF2-targeted compounds could be a potential option to monitor resistance and increase therapeutic efficacy of molecularly targeted alpha-therapy. Similarly, NTS-based radioligands have been developed for theranostic use ^44,45^. NTS/ NTS receptor signaling has, amongst others, been associated with prostate cancer exhibiting neuroendocrine features such as down-regulation or loss of AR and PSA ^46,47^. Thus, NTS-based imaging and therapy may be an option to monitor transition to to this very aggressive phenotype and to complement [^225^Ac]hu11B6 RIT. *Lastly, SCHLAP1* has been shown to be a reliable biomarker strongly associated with a shorter time to metastatic progression, biochemical relapse, and prostate-cancer-specific mortality ^48–50^.

The field of molecularly targeted alpha-therapy is currently undergoing clinical translation, including a planned [^225^Ac]hu11B6 trial. In the current study, we built on previous preclinical studies of hK2-targeted RIT by investigating novel methods to enhance therapeutic efficacy and explore prospective treatment strategies. To the best of our knowledge, this is the first study showing that FcRn-optimized IgG_3_ subforms can be applied to RIT. Although further studies are needed to replicate our findings, we also discovered a constellation of putative biomarkers associated with resistance to alpha-RIT.

## Methods

### Reagents and cell culture

Reagents were purchased from Sigma-Aldrich unless otherwise noted. Cell growth media were obtained from the Media Preparation Core Facility at Memorial Sloan Kettering Cancer Center (MSKCC, New York, NY). LNCaP-AR (LNCaP cell line with over-expression of wild-type AR and luciferase under the control of the ARR2-Pb promoter) was a kind gift from the laboratory of Dr. Charles Sawyers, which previously developed and reported the cell line ^51^. PC3 was purchased from American Type Culture Collection (ATCC). All cell lines were cultured according to the developer’s instructions and regularly tested for mycoplasma.

### Antibodies

Hu11B6-IgG_1_, hu11B6-IgG_3_^R435H^, hu11B6-IgG_3_, and hu11B6-IgG_1_^H435A^ used for an Actinium-225 labeling protocol using two steps were developed by DiaProst AB (Lund, Sweden) and produced by Innovagen AB (Lund, Sweden). Pre DOTA-conjugated hu11B6-IgG_1_ applicable for a one step labeling protocol was developed by Diaprost AB (Lund, Sweden) and produced by Fuji-Biosyth (London, England).

### Radiochemistry

Carrier-free solid ^225^Actiniumnitrate (5.80 × 10^4^ Ci/g; 214.6 × 10^4^ GBq/g), Oak Ridge National Laboratory, Oak Ridge, TN) was assayed at secular equilibrium with a CRC-15R radioisotope calibrator (Capintec, Inc., Florham Park, NJ). All reagents were American Chemical Society reagent grade or better. Radiolabeling procedures were performed using sterile metal-free and pyrogen-free plasticware (Corning, Inc.). The radiometal chelate 1,4,7,10-tetraazacyclododecane-1,4,7,10-tetraacetic acid (DOTA) was conjugated to the hu11B6 antibody to yield DOTA-hu11B6 and radiolabeled in one-step as previously described ^52^. Briefly, 0.005 mL of ^225^Ac-nitrate dissolved in 0.2 M HCl (Fisher Scientific) (0.1 mCi; 3.7 MBq) was mixed with 1.0 to 1.5 mg of DOTA-hu11B6-IgG_1_ and buffered with 0.1 mL of 2 M tetramethylammonium acetate and 0.01 mL of 150 g/L of L-ascorbic acid to pH 5.5. The clear and colorless reaction mixture was heated at 37°C for 75-110 minutes. The reaction was quenched with 0.02 mL of 50 mM diethylenetriaminepentaacetic acid and purified via size exclusion chromatography using a Bio-Rad 10DG column stationary phase with a 1% human serum albumin (HSA, Swiss Red Cross, Bern, Switzerland) and 0.9% sodium chloride (normal saline solution, Abbott Laboratories, North Chicago, IL) mobile phase. Four combinations of native and mutated, IgG_1_ and IgG_3_ isoforms of hu11B6 (i.e., native IgG_1_, H435A mutant IgG_1_, wild-type IgG_3_ and R435H mutant IgG_3_ were ^225^Ac-radiolabeled using a previously described two-step method to prepare alpha emitting nanogenerators ^53^. Radiochemical purity was determined by instant thin-layer chromatography using silica gel-impregnated paper (Gelman Science Inc., Ann Arbor, MI) using two mobile phases. Mobile phase I was 10 mM ethylenediaminetetraacetic acid; mobile phase II was 9% sodium chloride/10 mM sodium hydroxide. Activity was measured at secular equilibrium using a Packard gamma counter to count the signal in the 370-510 keV energy window. The binding kinetics of native [^225^Ac]DOTA-hu11B6, DOTA-hu11B6, and hu11B6 against recombinant hK2 analyte was measured by surface plasmon resonance (SPR) using a Biacore 2000 (GE Healthcare, Marlborough, MA) as previously described ^3^. Antibodies and constructs were captured on a Protein A sensor chip (GE Healthcare) ^54^. Antibody capture was performed by diluting each sample to a concentration of 0.001 mg/mL in HBS-EP buffer (BR100188, GE Healthcare), and flowing the solution over a protein A sensor chip for 1 min. at a flow rate of 0.005 mL/min. Binding kinetics of antibodies were evaluated across a concentration range of hK2 (0, 3.125, 6.25, 12.5, 25, 50, 100, and 200 nM) in HBS-EP buffer. The kinetic data was analyzed with Biacore 3.2.

### Animal studies

For xenograft studies, male athymic BALB/c nude mice (NU(NCr)-*Foxn1*^*nu*^; 6-8 weeks old, 20-25 g) were obtained from Charles River. LNCaP-AR tumors were induced in the left and right flanks by s.c. injection of 1-5 × 10^6^ cells in a 200 µL cell suspension of a 1:1 vol/vol mixture of medium with Matrigel (Collaborative Biomedical Products, Inc.). Tumors developed after approximately 3-7 weeks. The previously developed Hi-*Myc* x pb_*KLK2_* Hi-*Myc* GEMM, a prostate cancer-susceptible transgenic mouse model prostate tissue specific hK2 expression, was used in the studies ^12^. All interventions were performed under anesthesia (2% isoflurane).

### Pharmacokinetic tissue distribution

Biodistribution studies were conducted to investigate uptake and pharmacokinetic distribution of various Actinium-225 labeled IgG-subclasses (IgG_1_, IgG_1_^H435A^, IgG_3_, IgG_3_^R435H^) of hu11B6 ([^225^Ac]hu11B6) in the GEMM model, and LNCaP-AR and PC3 xenografts. Mice received a single 300 nCi (11 kBq) activity of [^225^Ac]hu11B6-IgG_1_, [^225^Ac]hu11B6^R435H^, [^225^Ac]hu11B6-IgG_1_, or [^225^Ac]hu11B6-IgG_1_^R435H^ (300 nCi on 5 μg antibody). Radioimmuno-compounds were administered via intravenous tail-vein injection (*t* = 0 hour). Animals (n=4–5 per group) were euthanized by CO_2_ asphyxiation at 4, 48, 120 and 360 h post-injection. Blood was immediately harvested by cardiac puncture. Tissues (including the tumor) were removed, rinsed in water, dried on paper, weighed, and counted on a gamma-counter using a 370–510 keV energy window at secular equilibrium. Aliquots (0.020 mL) of the injected activities were used as decay correction standards and background signal was subtracted from each sample. The %IA/g was calculated for each animal and data plotted as mean±SD.

### Therapy studies

In the initial studies, the therapeutic efficacies of 300 nCi (11 kBq) [^225^Ac] on 5 μg hu11B6-IgG_1_, hu11B6-IgG_1_^H435A^, hu11B6-IgG_3_, and hu11B6IgG_3_^R435H^, labeled using a two-step protocol, were compared in s.c. LNCaP-AR tumors and GEMM. In a subsequent study, two different treatment regimens of labeled hu11B6-IgG_1_ labeled using a one-step protocol were evaluated; single activity treatment (600 nCi; 22 kBq) was compared to fractionated administration (300 + 300 nCi; 11 + 11 kBq) [^225^Ac]hu11B6-IgG_1_ in s.c. LNCaP-AR tumors (5 mice/treatment group and 5 untreated mice/treatment group). Length (l) and width (w) of the tumors were measured by caliper and tumor volume was calculated using the modified formula for a rotated ellipsoid (*V*=½ w^2^l) ^55^ was calculated. Weight loss of 20% or a tumor diameter exceeding 15 mm was set as an endpoint. Survival percentages over time were calculated to study therapy effectiveness in both studies.

### Measurement of kallikreins

Free (fPSA) and total (tPSA) PSA were determined with the dual-label DELFIA immunofluorometric assay (Prostatus™ PSA Free/Total PSA from Perkin-Elmer Life Sciences Turku, Finland). This assay determines free and complexed PSA in an equimolar fashion, and the cross-reaction for PSA-alpha 1-antichymotrypsin (PSA-ACT) in the fPSA assay is below 0.2% ^56^. The lower limit of detection for tPSA is 0.10 ng/ml (coefficient of variation of 5.0% at 2.32 ng/mL) and for fPSA 0.04 ng/mL (coefficient of variation of 5.9% at 0.25 ng/mL). hK2 was measured using an in-house research assay protocol ^57^; the detection limit of the assay is 0.005 ng/mL with assay imprecision values (mean coefficients of variation) ranging from 5.7% to 11% for high and low hK2 controls, respectively.

### RNA extraction

RNA from snap frozen LNCaP-AR tumors was extracted using the RNeasy Mini Kit (QIAGEN) according to the instructions provided by the manufacturer.

### Transcriptome sequencing (RNA sequencing)

After RiboGreen quantification and quality control by Agilent BioAnalyzer, 500 ng of total RNA underwent polyA selection and TruSeq library preparation according to instructions provided by Illumina (RS-122-2102, TruSeq Stranded mRNA LT Kit) with 8 cycles of PCR. Samples were barcoded and run on a HiSeq 4000 in a 50bp/50bp paired end run, using the HiSeq 3000/4000 SBS Kit (Illumina). An average of 44 million paired reads was generated per sample. Ribosomal reads represented maximally 1.2% of the reads generated and the percent of mRNA bases averaged 63%.

### Transcriptome sequencing analysis

Raw read count RNA-sequencing gene expression data for a total of 58,828 genes were acquired from tumor samples treated with 2x 300nCi (2x 11 kBq) [^225^Ac]hu11B6-IgG_1_ (n=3) and left untreated (n=3). After confirming the quality of sequences via FastQC, principal component analysis (PCA) was performed in R, where the PCA plot demonstrated a clear division between the samples from the two cohorts (**Supplemental Figure 3**). Hierarchical clustering of samples using the Manhattan method for distance calculation further demonstrated a clear divide between the two cohorts. To visualize the magnitude of effect and to highlight the groups of genes that were similar to each other based on their expression, a heatmap was plotted based on the log2 raw read counts (**Supplemental Figure 4**).

Differential expression analysis of RNA-sequencing expression profiles was performed using Bioconductor’s edgeR^58^ package in R. edgeR implements statistical methods based on negative binomial distributions, namely empirical Bayes methods, that permit the estimation of gene-specific biological variation, even for experiments with minimal levels of biological replication. Raw read counts are provided as input where edgeR introduces possible bias sources into the statistical model to perform an integrated normalization followed by differential expression analysis. P-values and fold change in log 2 were calculated for each gene. P-values were adjusted for False Discovery Rate (FDR) using the Benjamini-Hochberg method. Differentially expressed genes were then selected based on the following criteria: Adjusted p-value <0.001 and absolute value of log2 fold change >2. A positive fold change represented up-regulation, whereas a negative fold change represented down-regulation in tumor samples treated with [^225^Ac]hu11B6-IgG^1^.

### Statistics

All statistical analysis of data, except for analysis of RNA-sequencing data (see paragraph above), was performed using Prism software (Graphpad Software Inc, La Jolla, CA). Data are represented as mean±SD or median and range (survival data) (n≥3 independent samples/group for all analyses). No datapoints were excluded from analysis. Unpaired, two-tailed t-tests were used for comparison of two groups. Analysis of survival data was performed by the log-rank test (Mantel-Cox). A p-value<0.05 was considered to indicate statistical significant differences.

### Ethical approval

All animal experiments were conducted in compliance with institutional Animal Care and Use Committee (IACUC)-established guidelines at MSKCC and under supervision by the MSKCC Research Animal Resource Center.

## Author Contributions

### Conception and design

David Ulmert, Robert J. Klein

### Development of methodology

Sven-Erik Strand, Hans Lilja, Christin Malmborg Hager, Michael R McDevitt, Robert J. Klein, David Ulmert

### Acquisition of data (provided animals, acquired and managed patients, provided facilities, etc.)

Steven M Larson, Hans Lilja, David Ulmert, Michael R McDevitt, Robert J. Klein

### Analysis and interpretation of data (e.g., statistical analysis, biostatistics, computational analysis)

Mesude Bicak, Katharina Lückerath, Michael R McDevitt, Robert J. Klein, David Ulmert

### Writing, review, and/or revision of the manuscript

Mesude Bicak, Katharina Lückerath, Teja Kalidindi, Sven-Erik Strand, Michael Morris, Caius Radu, Robert Damoiseaux, Norbert Peekhaus, Austin Ho, Darren Veach, Ann-Christin Malmborg Hager, Steven M Larson, Hans Lilja, Michael R McDevitt, Robert J. Klein, David Ulmert

### Administrative, technical, or material support (i.e., reporting or organizing data, constructing databases)

Mesude Bicak, Katharina Lückerath, Robert J. Klein, David Ulmert

### Study supervision

Steven M Larson, Michael R McDevitt, Robert J. Klein, David Ulmert

## Notes

Conflict of interests: Sven-Erik Strand, Ann-Christin Malmborg-Hager, Hans Lilja and David Ulmert are consultant/advisory board members for and hold ownership interest in Diaprost AB. Sven-Erik Strand and David Ulmert are listed as co-inventors on several patents regarding the humanized form of 11B6, which is owned by Diaprost. Caius Radu is co-founder and holds equity in Sofie Biosciences and Trethera Therapeutics. Intellectual property has been patented by the University of California and has been licensed to Sofie Biosciences and Trethera Therapeutics. This intellectual property was not used in the current study. Hans Lilja has patents on assays to measure intact PSA and a statistical method to detect prostate cancer commercialized as 4Kscore by OPKO Health, receives royalties from sales of this test, and owns stock in OPKO Health. Steven M. Larson reports receiving commercial research grants from Regeneron and Telix, holds ownership interest (including patents) in Voreyda, Imaginab, and Elucida, and is a consultant/advisory board member for Johnson and Johnson. Memorial Sloan Kettering Cancer Center has filed for IP protection for inventions related to alpha particle technology of which Michael McDevitt is an inventor. Michael McDevitt was a consultant for Actinium Pharmaceuticals, Regeneron, Progenics, Bridge Medicines, and General Electric. No other potential conflict of interest relevant to this article was reported.

