## Supplemental Figures 1-4 for "Genetic signature of prostate cancer resistant to optimized hK2 targeted alpha-particle therapy"

### Supplemental Data

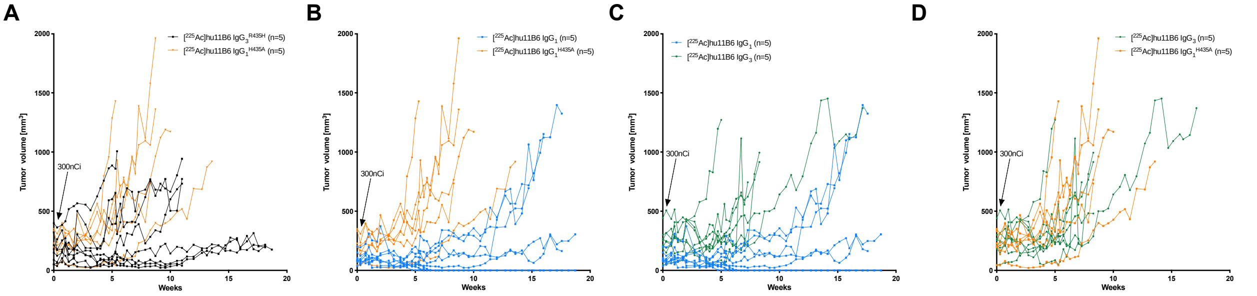


**Supplementary Figure 1. The impact of IgG subclass on [$^{225}$Ac]hu11B6 alpha-radioimmunotherapy.** Mice with LNCaP-AR s.c. xenografts were randomized to receive 300 nCi (11 kBq) [$^{225}$Ac]hu11B6-IgG$_1$ (n=5; blue),[$^{225}$Ac]hu11B6-IgG$_1^{H435A}$ (n=5; orange), [$^{225}$Ac]hu11B6-IgG$_3^{R435H}$ (n=5; black) or [$^{225}$Ac]hu11B6-IgG$_3$ (n=5; green). Tumor volumes in individual mice following treatment with IgG$_3^{R435H}$ vs. IgG$_1^{H435A}$ (**A**), IgG$_1$ vs. IgG$_1^{H435A}$ (**B**), IgG$_1$ vs. IgG$_3$ (**C**), or IgG$_3$ vs. IgG$_1^{H435A}$ (**D**) antibodies.

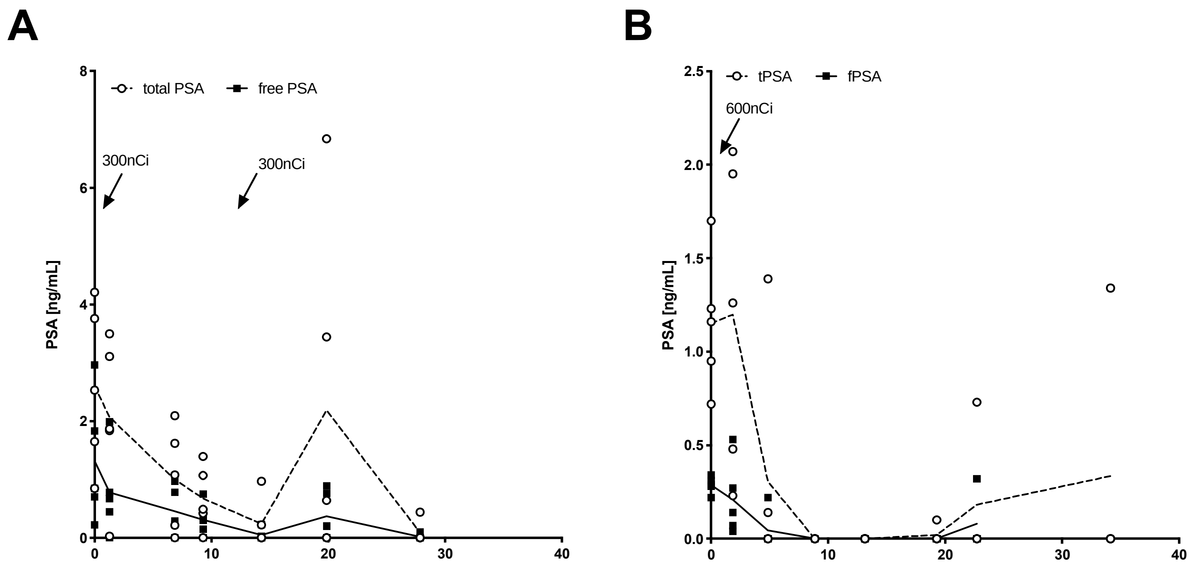


**Supplementary Figure 2. Therapeutic efficacy of fractionated vs. single high-activity [$^{225}$Ac]hu11B6-IgG$_1$ alpha RIT.** Mice with LNCaP-AR s.c. xenografts received 2x 300nCi (2x 11 kBq; ~4.5 month apart) (**A**) or 1x 600 nCi (22 kBq) (**B**) [$^{225}$Ac]hu11B6-IgG$_1$. Total (open circles, dotted line) and free (filled circles, solid line) PSA was measured at indicated times as surrogate marker for tumor burden (n=5 mice/group). Note the different scales on the y-axes.

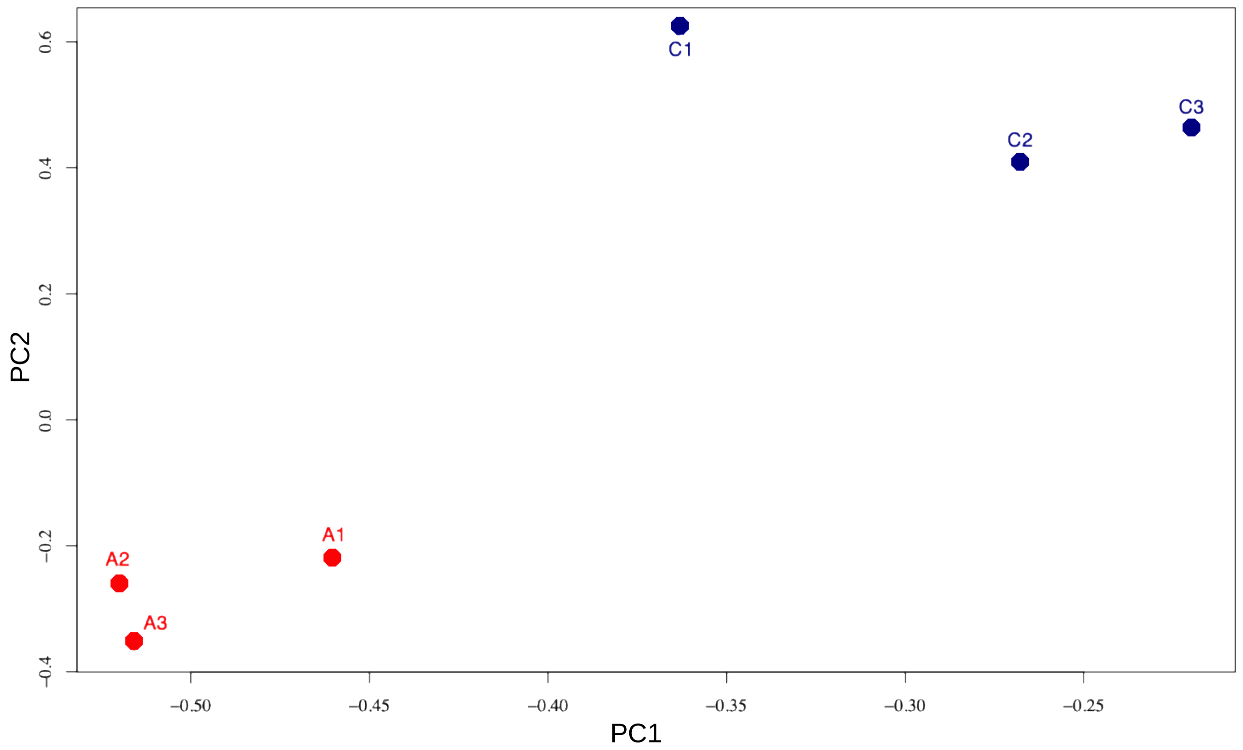


**Supplemental Figure 3. Principal component analysis (PCA).** PCA of tumors treated with [$^{225}$Ac]hu11B6-IgG$_1$ (red) vs. untreated control samples (blue). The x-axis represents PC1, while the y-axis represents PC2. Samples that have similar expression profiles are clustered together. The PCA shows that the two experimental groups are distinctively separated along both PCs.

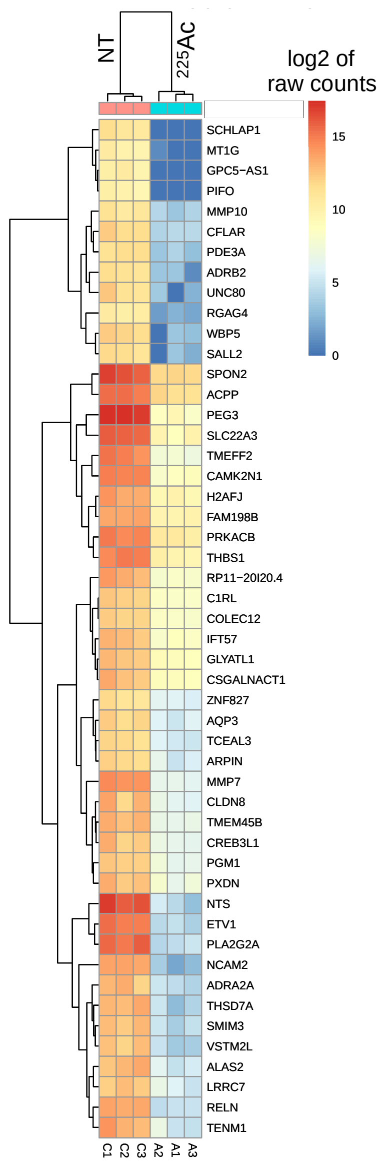


**Supplemental Figure 4. Heatmap of the top 50 differentially expressed genes.** The heatmap displays log2 values of raw readcounts for top 50 DEGs. A higher readcount indicates a higher expression of a particular gene for each sample. Both raw (pre-analysis) and processed (post-analysis) data can be used to plot heatmaps. Here we chose to plot a heatmap of raw gene expression for the top 50 DEGs to demonstrate the similarity within treated and untreated samples, since treated samples for all top 50 DEGs have a lower raw gene expression than untreated samples. Furthermore, as indicated by the color scheme, when it is closer to the red shade versus blue shade, this represents a more significant difference per gene between the two groups (i.e *NTS*, *ETV1*, *NCAM2* clustered closer together compared to *SCHLAP1* even among top 50 DGEs themselves), which is inline with the findings from our DEG analysis (**Figure 5**). Samples from tumors treated with [$^{225}$Ac]hu11B6-IgG$_1$ are shown in the three columns on the right (A1, A2, A3); samples from untreated tumors are shown in the left three columns (C1, C2, C3). *$^{225}$Ac - [$^{225}$Ac]hu11B6-IgG$_1$ treated tumors; NT – not treated tumors.*
